# Neural Responses and Perceptual Sensitivity to Sound Depend on Sound-Level Statistics

**DOI:** 10.1101/850339

**Authors:** Björn Herrmann, Thomas Augereau, Ingrid S. Johnsrude

**Author notes:** Correspondence concerning this article should be addressed to Björn Herrmann, Rotman Research Institute, Baycrest, 3560 Bathurst St, North York, ON, M6A 2E1, Canada.

## Abstract

Sensitivity to sound-level statistics is crucial for optimal perception, but research has focused mostly on neurophysiological recordings, whereas behavioral evidence is sparse. We use electroencephalography (EEG) and behavioral methods to investigate how sound-level statistics affect neural activity and the detection of near-threshold changes in sound amplitude. We presented noise bursts with sound levels drawn from distributions with either a low or a high modal sound level. One participant group listened to the stimulation while EEG was recorded (Experiment I). A second group performed a behavioral amplitude-modulation detection task (Experiment II). Neural activity depended on sound-level statistical context in two different ways. Consistent with an account positing that the sensitivity of neurons to sound intensity adapts to ambient sound level, responses for higher-intensity bursts were larger in low-mode than high-mode contexts, whereas responses for lower-intensity bursts did not differ between contexts. In contrast, a concurrent slow neural response indicated prediction-error processing: The response was larger for bursts at intensities that deviated from the predicted statistical context compared to those not deviating. Behavioral responses were consistent with prediction-error processing, but not with neural adaptation. Hence, neural activity adapts to sound-level statistics, but fine-tuning of perceptual sensitivity appears to involve neural prediction-error responses.

## Introduction

Acoustic environments are highly variable. In the course of a day, someone may work in a library, commute by subway, listen to a podcast, and attend a stadium concert. How does that person manage to hear effectively in all those different environments? Some perceptual flexibility is crucial for the optimization of perception and behavior^1–4^. However, neurons that support audition are limited in the range of sensory stimulation to which they respond^5,6^. For example, spiking activity of auditory nerve fibers in rodents sensitively discriminates between different sound levels over a limited range, typically between 10 and 40 decibels^7,8^. Sound levels outside this range either saturate neural responding (i.e., when sounds are more intense than the upper response limit) or do not elicit a response (i.e., when sounds are less intense than the lower response limit; Figure 1). Humans can perceptually identify small differences in the levels of two sounds over a 120-decibels range, even though individual neurons only respond over a ~40 dB range^6,9^.

**Figure 1:**
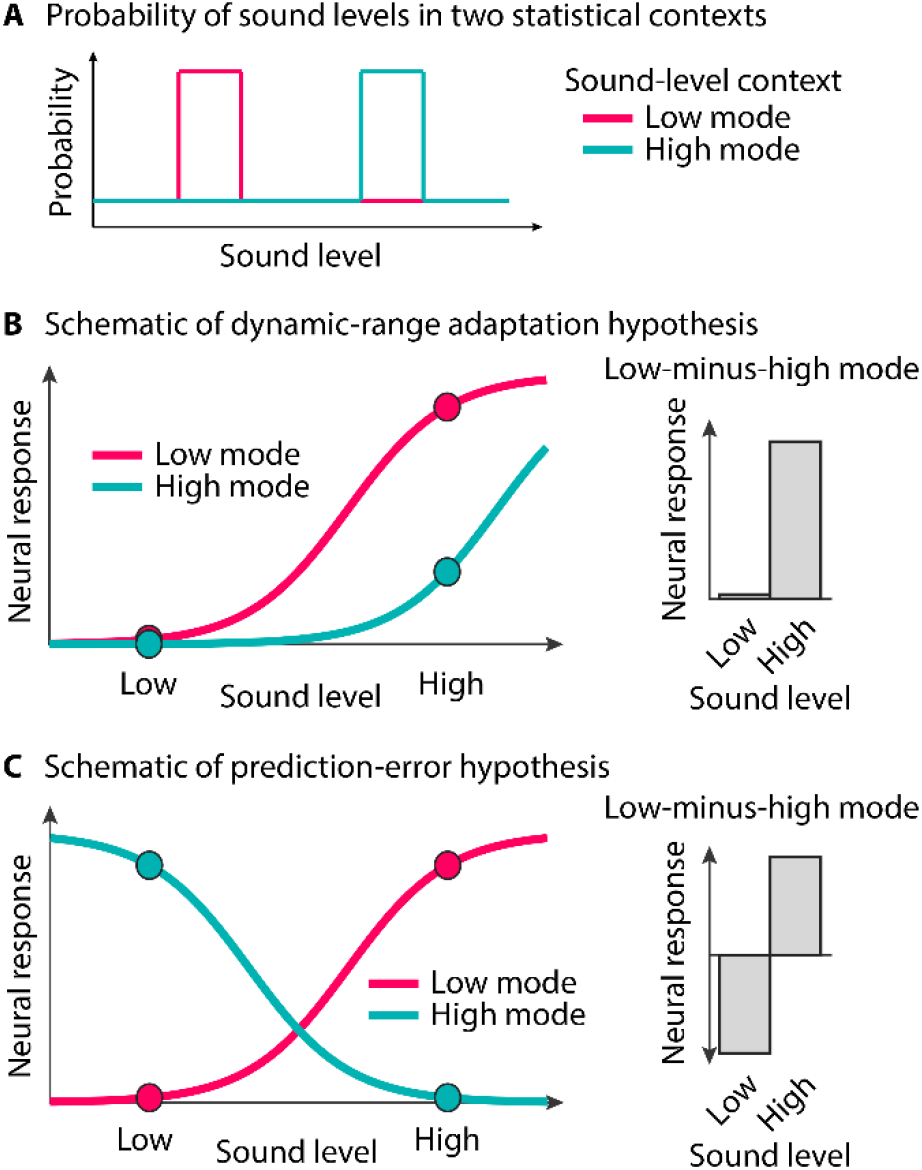
Schematic depiction of hypothesized effects of sound-level statistics on neural responses and behavior. **A:** Probability with which sound levels occur in two statistical contexts **B:** Neural responses as a function of sound level for two sound-level contexts with different modal sound levels (low vs. high; panel A) predicted by the dynamic-range adaptation hypothesis. Dots mark two target-intensities (lower and higher), similar to the neural and behavioral analysis of empirical data (Experiment I & II). Gray bars indicate the difference between the low-mode and the high-mode contexts, separately for each target intensity^10,11,21^. **C:** Similar as in panel B for the prediction-error hypothesis.

Achieving a wide perceptual range with a limited neuronal range is thought to be accomplished by a dynamic adjustment of the response range (input-output function) of neurons to sound-level statistics in the environment (Figure 1A)^10,11^. This process - called adaptation to stimulus statistics, dynamic-range adaptation, or gain control^1,12–16^ - involves shifting the response range of neurons depending on the mean or modal sound level in an acoustic environment^10,11,17–22^. When ambient sound levels are low, neurons are more sensitive and more responsive, but saturate for moderate-to high-level sounds, compared to when ambient sound levels are moderate to high (red and teal lines in Figure 1A,B, respectively). The red and teal dots in Figure 1B indicate predicted responses to target stimuli of lower or higher intensity, presented in two different sound contexts (low ambient level; high ambient level). According to this dynamic-range adaptation hypothesis, if adaptation shifts the neural response range based on sound-level statistical context, we should observe a large effect of context on a higher-level target sound, and much less effect of context on a lower-level sound (Figure 1B, right).

A different line of work indicates that neural response to sound may reflect the deviation of sound level from predicted level^23–26^. When ambient sound levels are low, neurons are more responsive to higher-level than lower-level sounds, but when ambient sound level are high, neurons are more responsive to lower-level than higher-level sounds^23,25^. The red and teal dots in Figure 1C indicate the predicted responses to target stimuli of lower and higher intensity, presented in two different sound contexts. Under this scenario, we should observe a large effect of context for low-level and high-level target sounds, albeit of opposite direction (Figure 1C, right). Effects of this nature may reflect an error signal when a sound deviates from the predicted statistical context^27–30^.

Most research on adaptation of neural activity to sound-level statistics has been conducted using neurophysiological techniques in non-human mammals^10,11,13,17–20,31,32^. We have recently demonstrated using magnetoencephalography that neural activity in human auditory cortex adapts to sound-level statistics (i.e., to the modal sound-level) consistent with dynamic-range adaptation (Figure 1B)^21^. However, whether such adaptive changes confer benefit to perceptual sensitivity is less clear. We expect that the ability to perceive changes in sound level, such as those introduced by amplitude modulation, will also be affected by sound-level statistical context (e.g., low vs. high mode), in a way that reflects the neurophysiological data: Dynamic-range adaptation predicts better sensitivity for higher-level target sounds than lower-level target sounds, uniquely in the context with a low modal level (Figure 1B, right). A prediction-error account predicts better sensitivity for target sound levels that deviate from a distribution’s modal sound level; that is, better sensitivity for higher-level sounds in low-mode contexts and better for sensitivity lower-level sounds in high-mode contexts (Figure 1C, right).

In a behavioral study, Simpson and colleagues^33^ assessed whether relative loudness perception depends on the modal sound level of the auditory context. Participants were presented with sound pairs whose sound levels were drawn from a distribution with either a low or a high modal level, and they indicated which of the two sounds was louder. Discrimination performance was sensitive to the context’s modal level. However, inconsistent with dynamic-range adaptation measured neurophysiologically (Figure 1B)^10,11,17,18,21^, sound-level discrimination was better for low-intensity compared to higher-intensity target sounds in the high-modal context^33^. Neurophysiological work on dynamic-range adaptation, in contrast, suggests that neurons are not very sensitive to low-intensity sounds when the ambient sound level is high (Figure 1B, right). The reported behavioral results are more consistent with prediction-error responses (Figure 1C). The experimental paradigm used in this previous study^33^ differed in a potentially important way from that typically used to study neural adaptation to sound-level statistics^10,11,17,18,31^: the interval between sound pairs was substantially longer to allow for a behavioral response^33^. It is not clear whether neural adaptation to sound-level statistics is effective over these longer intervals.

In the current study, we use a common stimulation paradigm to examine the effects of sound-level statistics on neural responses (Experiment I) and perceptual sensitivity to fluctuation in a sound’s amplitude (Experiment II). Young adults listened to white-noise bursts (0.1 s duration; 0.5 s onset-to-onset interval) whose sound levels were drawn from a distribution with either a 15-dB or a 45-dB modal level (Figure 2). The main analyses focused on neural and behavioral responses to noise bursts (targets) for which the sound level, and the sound level of the directly preceding noise, were identical across the two statistical contexts (black dots in Figure 2, middle/right). This allowed us to investigate the effects of longer-term sound-level statistics on neural responses and perception^21^.

**Figure 2:**
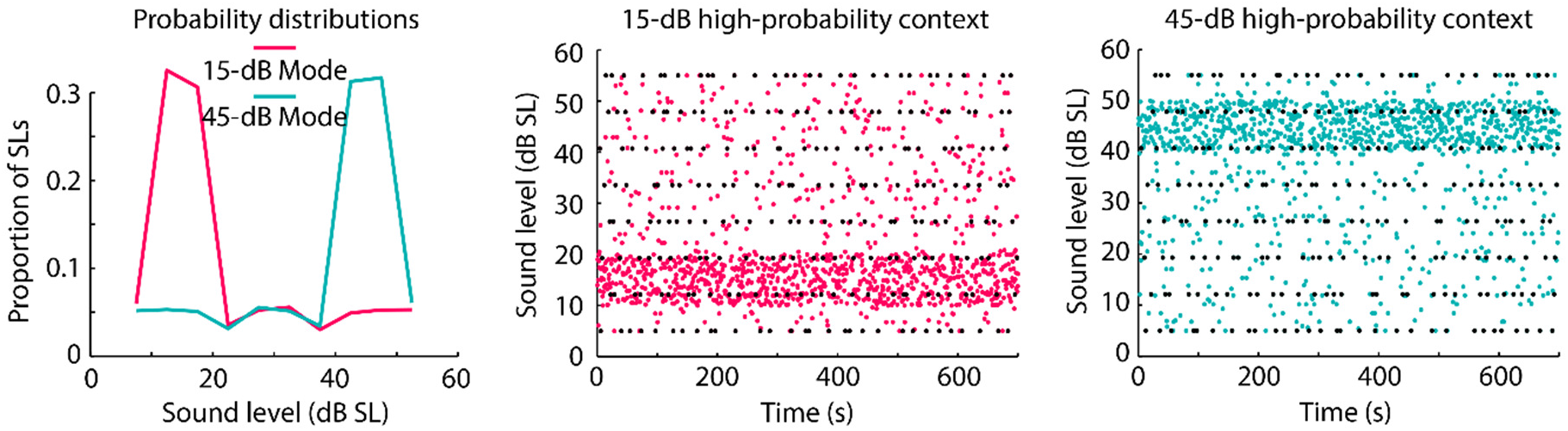
Experimental stimulation used in Experiment I and II. **Left:** Example probability distributions used for the acoustic stimulation for the two statistical contexts: one had a modal sound level of 15 dB SL, the other one had a modal sound level of 45 dB SL. **Middle/right:** 1400 white noises with different sound levels (y-axis) were presented at a rate of 0.5 s within a 700-s block (x-axis). Each dot reflects the sound level of one noise stimulus. Black dots indicate the stimuli of interest (targets) for which the effect of sound-level statistics on neural responses (Experiment I) and amplitude-modulation detection performance (Experiment II) was analyzed. For each target burst, its level and the level of the directly preceding noise burst were identical across contexts (15 dB SL and 45 dB SL modal level).

## Results

### Experiment I: Neural responses depend on sound intensity and sound-level context

Electroencephalography (EEG) was recorded from twenty-five normal-hearing adults (18-31 years), while they listened to 0.1-s white-noise bursts presented in two sound-level contexts (Figure 2). Participants watched a muted, subtitled movie during the experiment. Sensitivity of neural responses to sound level was assessed by binning the responses to each sound (across both contexts) into five sound-level categories (20-dB width, centered on 15, 22.5, 30, 37.5, and 45 dB SL^21^). Single-trial responses within each category were averaged (Figure 3A) and the P1-N1 and P2-N1 peak-to-peak amplitudes were calculated separately for each participant^34–38^. A linear function was fit to amplitude data (separately for P1-N1 and P2-N1) as a function of the five sound levels, independently for each participant. Slopes (linear coefficients) were reliably larger than zero for the P2-N1 amplitude (t_24_ = 4.044, p = 4.7×10^−4^, r_e_ = 0.637; Figure 3C), demonstrating that the P2-N1 amplitude increased with increasing sound level ^39–41^. The slope for the P1-N1 amplitude, while numerically positive, did not differ from zero (t_24_ = 1.236, p = 0.229, r_e_ = 0.245; Figure 3B).

**Figure 3:**
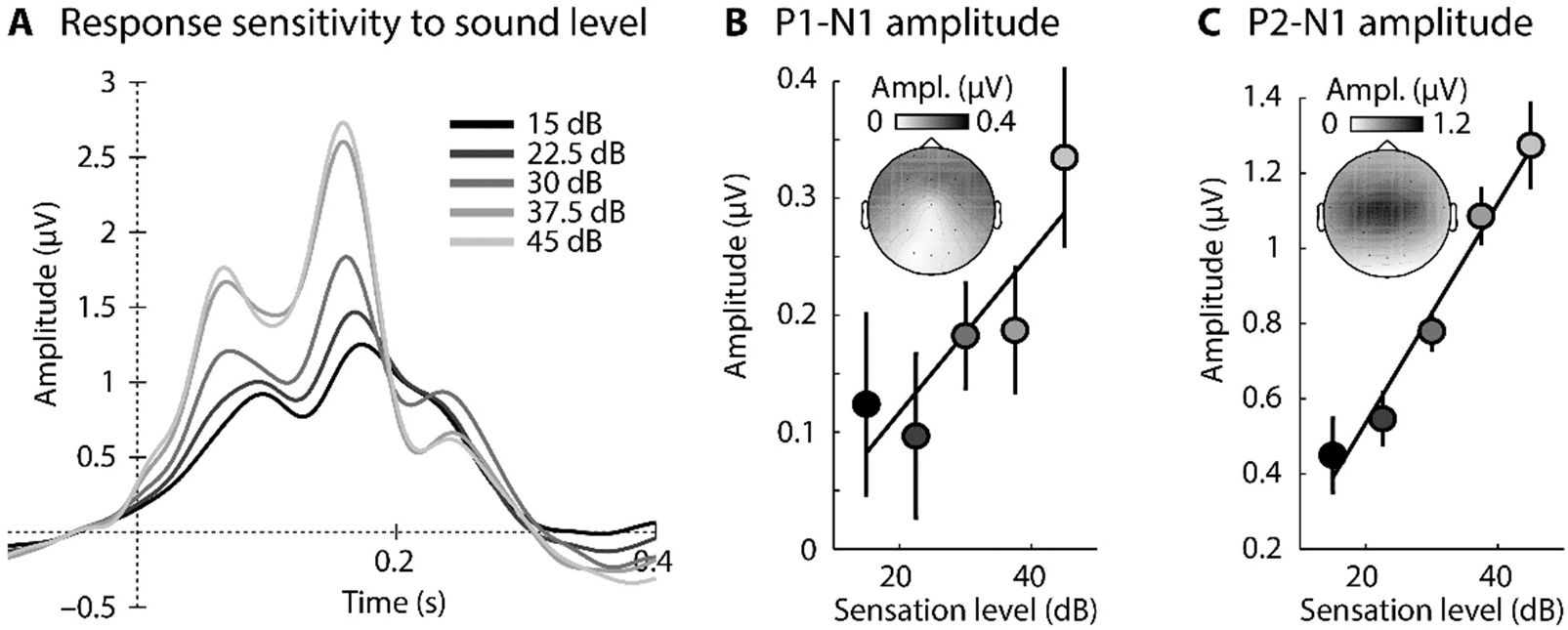
Overall response sensitivity to sound level (across sound-level contexts). **A:** Response time courses for five different sound-level categories (averaged across a fronto-central electrode cluster). **B:** P1-N1 peak-to-peak amplitude for each sound level. The solid line reflects the averaged slope of the linear fit. Error bars reflect the standard error of the mean (removal of between-subject variance^42^). **C:** Same as in panel B for P2-N1 peak-to-peak amplitude.

P1-N1 and P2-N1 responses to noise bursts were compared between two different sound-level contexts presented in three blocks each: the modal level of the noise bursts was 15 dB in one context and 45 dB in the other (Figure 2)^21^. Critically, analyses focused on noise bursts for which the intensity, and the intensity of the directly preceding burst, were identical across contexts (these targets are indicated by black dots in Figure 2, middle/right). Hence, any difference in neural response between stimuli from the different contexts must be due to the longer-term sound-level distributions.

In order to maximize the signal-to-noise ratio of the neural responses to target bursts, they were binned according to target intensity, separately for the two statistical contexts: responses from targets with intensities below 30 dB SL were combined (mean of 15.7 dB SL), as were those with intensities higher than 30 dB SL (mean of 44.3 dB SL). The P1-N1 and P2-N1 peak-to-peak amplitudes were calculated for both contexts, for both target-intensity categories, separately for each participant (Figure 4B).

**Figure 4:**
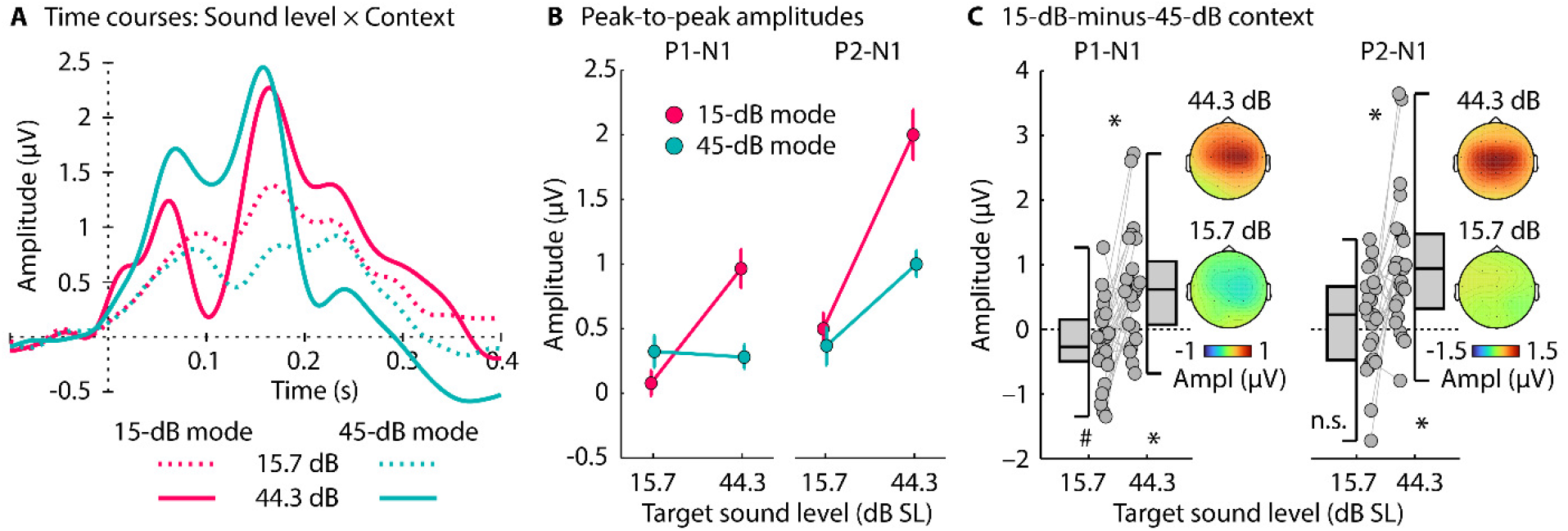
Neural responses to noise bursts in two different sound-level contexts. **A:** Response time courses (averaged across a fronto-central electrode cluster) for sound level × context conditions. **B:** P1-N1 and P2-N1 peak-to-peak amplitudes for target-noise bursts with different intensities (one centered on 15.7 dB and one on 44.3 dB) for the 15-dB SL context and for the 45-dB SL context. Error bars reflect the standard error of the mean (removal of between-subject variance^42^). The interaction between context and target-intensity category is significant for both P1-N1 and P2-N1 amplitudes (p < 0.05). **C:** Amplitude difference between the 15-dB SL and 45-dB SL contexts, separately for low- and high-intensity targets. A positive value means a larger amplitude for the 15-dB SL compared to the 45-dB SL context, whereas a negative value means a larger amplitude for the 45-dB SL compared to the 15-dB SL context. Topographical distributions reflect the mean response difference between the two statistical contexts. #p ≤ 0.01, *p ≤ 0.05, n.s. - not significant

A repeated-measures analysis of variance (rmANOVA) revealed that the P1-N1 amplitude increased from the low to high Target-Intensity Category (F_1,24_ = 5.446, p = 0.028, η^2^_p_ = 0.185), and was larger for the 15-dB SL compared to the 45-dB SL context (F_1,24_ = 4.485, p = 0.045, η^2^_p_ = 0.158). The Context × Target-Intensity Category interaction was significant (F_1,24_ = 18.854, p = 2.2×10^−4^, η^2^_p_ = 0.440): P1-N1 amplitudes were larger in the 15-dB SL compared to the 45-dB SL context for the higher-intensity target category (44.3 dB; t_24_ = 4.021, p = 0.0005, r_e_ = 0.635), but not for the lower-intensity target category, for which we observed a trend towards larger responses in the 45-dB SL compared to the 15-dB SL context (15.7 dB; t_24_ = −1.979, p = 0.059, r_e_ = 0.375; Figure 4C).

The P2-N1 amplitude also increased from the low to high Target-Intensity Category (F_1,24_ = 23.730, p = 5.8×10^−5^, η^2^_p_ = 0.497), and was larger for the 15-dB SL compared to the 45-dB SL context (F_1,24_ = 21.632, p = 0.0001, η^2^_p_ = 0.474). The Context × Target-Intensity Category interaction was significant (F_1,24_ = 9.923, p = 0.0043, η^2^_p_ = 0.293): P2-N1 amplitudes were larger in the 15-dB SL compared to the 45-dB SL context for the higher-intensity target category (44.3 dB; t_24_ = 4.717, p = 8.5×10^−5^, r_e_ = 0.694), whereas no difference between contexts was observed for the lower-intensity target category (15.7 dB; t_24_ = 0.901, p = 0.377, r_e_ = 0.181; Figure 4C; compare to predictions in Figure 1B, right). Topographical distributions suggest that auditory cortex underlies the observed P1-N1 and P2-N1 neural responses (Figure 4C)^43,44^.

The results are consistent with recent work in humans using sine tones^21^ and with previous work in animals^10,11,17,18^, and are largely consistent with a dynamic-range adaptation account (Figure 1C), particularly for P2-N1 responses. The data suggest that response sensitivity shifts dynamically with modal level, enhancing sensitivity to higher-level sounds in contexts with a low modal level (see Figure 1C). However, time courses in Figure 4A also suggest that a slower, context-dependent neural response may be superimposed on the P1-N1-P2 responses. In order to isolate the slow neural activity, response time courses for the two statistical contexts were subtracted (15-dB mode minus 45-dB mode), separately for the two target-intensity categories (15.7 dB, 44.3 dB; Figure 5).

**Figure 5:**
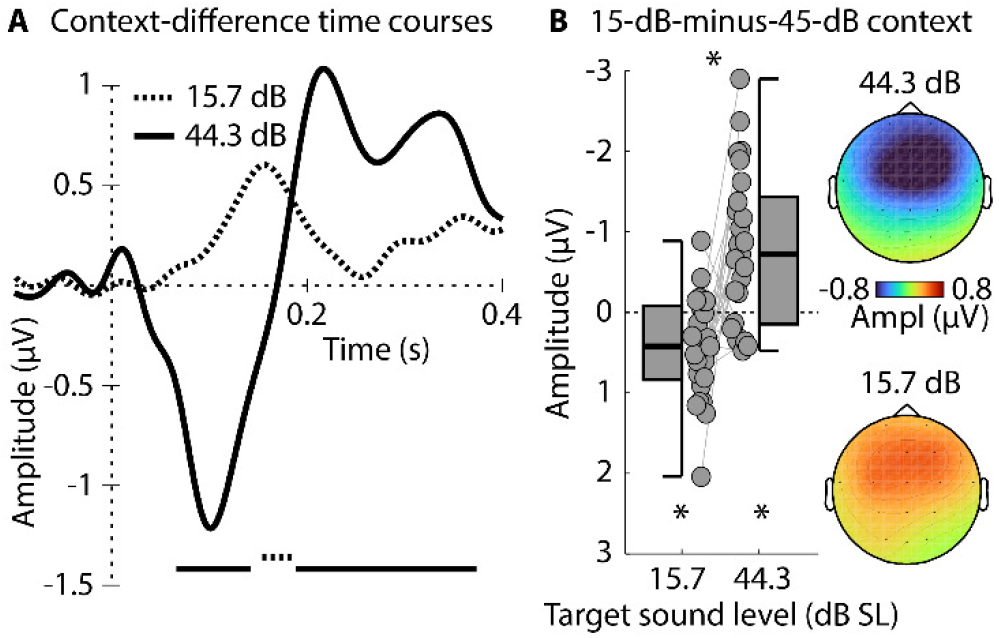
Response difference between the two statistical contexts (15-dB mode minus 45-dB mode). **A:** Difference time courses for the two target-intensity categories (15.7 dB, 44.3 dB). The lines at the bottom reflect a significant difference from zero (FDR-corrected^45,46^). **B:** Amplitude difference between the 15-dB SL and 45-dB SL contexts, separately for low- and high-intensity targets (time windows 0.1-0.2 s and 0.05-0.15 s, respectively). Note that negative amplitude differences are plotted upwards, in order to facilitate correspondence with Figures 1, 4, and 7. The reversed polarity relative to the other figures is the result of the negative potential, possibly reflecting a prediction-error signal. Topographical distributions reflect the mean response difference between the two statistical contexts. *p ≤ 0.05

Consistent with a slow negative potential reflecting a prediction-error signal (Figure 1C), responses were more negative for higher- than lower-intensity targets in the 15-dB SL context (0.05-0.15 s; t_24_ = −4.154, p = 3.6×10^−4^, r_e_ = 0.647; Figure 5A, negative-going wave) and more negative for lower- than higher-intensity targets in the 45-dB SL context (0.1-0.2 s; t_24_ = 3.496, p = 0.0019, r_e_ = 0.581; Figure 5A, positive-going wave), reflecting a Context × Target-Intensity Category interaction (t_24_ = 4.582, p = 1.2×10^−4^, r_e_ = 0.683; compare Figure 5B to predictions in Figure 1C, right). Topographical distributions (Figure 5B) are consistent with underlying sources in auditory cortex^43,44^. The results from the analysis of the slow negative potential mirror previous observations of a mismatch negativity elicited when a sound level deviates from a predicted level^23–26^.

In sum, the results of Experiment I show that P1-N1 and P2-N1 responses generated in auditory cortex are largely consistent with a dynamic-range adaptation hypothesis (Figure 1B), but that a slower negative-going response elicited concurrently indexes a prediction-error response (Figure 1C). In Experiment II, we used to same paradigm as in Experiment I in order to elucidate the perceptual consequences of such neural changes.

### Experiment II: Perceptual sensitivity to AM depends on sound-level context

The effect of sound-level statistics (context) on perceptual sensitivity was investigated by applying amplitude modulation (AM; 50 Hz) to the target sounds of Experiment I and comparing detection performance across contexts (non-target, filler, noise bursts were unmodulated). An AM-detection task was chosen because AMs are fluctuations in sound level, and neural adaptation to sound-level statistics is hypothesized to affect sound-level acuity.

In order to equate the difficulty of AM detection across the target-sound levels, six participants (20-27 years) who did not participate in Experiment I or II participated in a pre-test to estimate the function (linear slope) that relates AM detection thresholds to target-sound levels. This function was then applied to the AM detection threshold that was estimated for one sound level (40 dB SL) for each participant of Experiment II (N = 33; aged 17-31 years; all participants were naive), in order to determine the AM depth that yields 75% detected targets for each of the eight target intensity levels of the experimental protocol depicted in Figure 2 (see Experiment II pre-test methods and Figure 6, for details).

**Figure 6:**
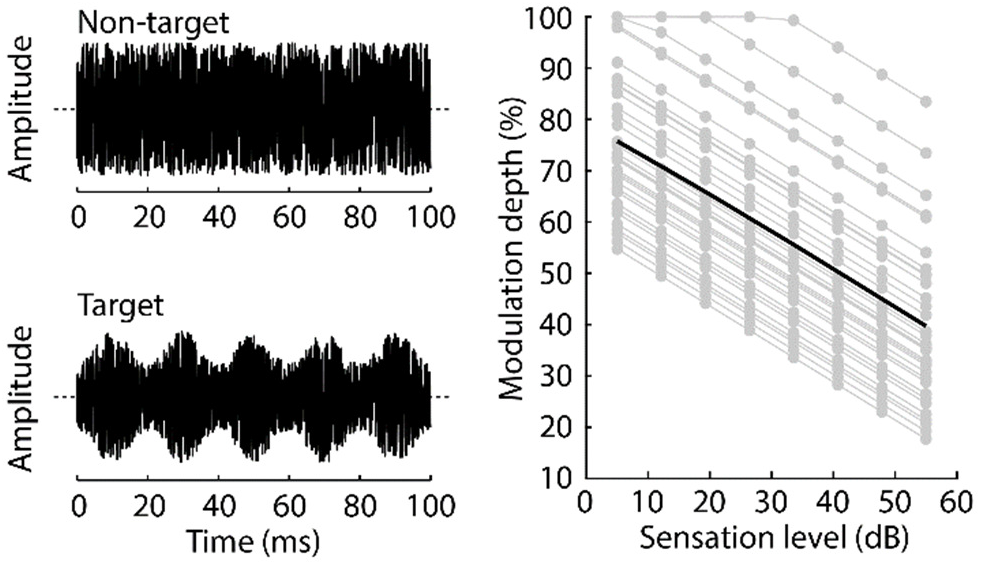
Acoustic stimuli and AM depths for behavioral Experiment II. **Left:** Example waveforms of non-target (filler, unmodulated) and target (50-Hz amplitude-modulated) noise bursts used in Experiment II. In the example, the amplitude of the target noise burst is modulated with a modulation depth of 50%. **Right:** AM depths used for each participant (gray lines) and sound level (gray dots) in Experiment II. The black line reflects the mean across participants. The slope common to all lines was calculated from the Experiment II pre-test. The height of each participant’s line is determined by the AM depth yielding 75% correct detection in 40 dB SL noise bursts during preliminary testing.

As in Experiment I, responses to targets were binned according to target intensity, separately for the two contexts: responses from targets with an intensity below 30 dB SL (mean of 15.7 dB SL) were combined, as were those with an intensity higher than 30 dB SL (mean of 44.3 dB SL). Perceptual sensitivity (d’) was calculated for both target-intensity categories and for both statistical contexts (Figure 7A), for each participant.

**Figure 7:**
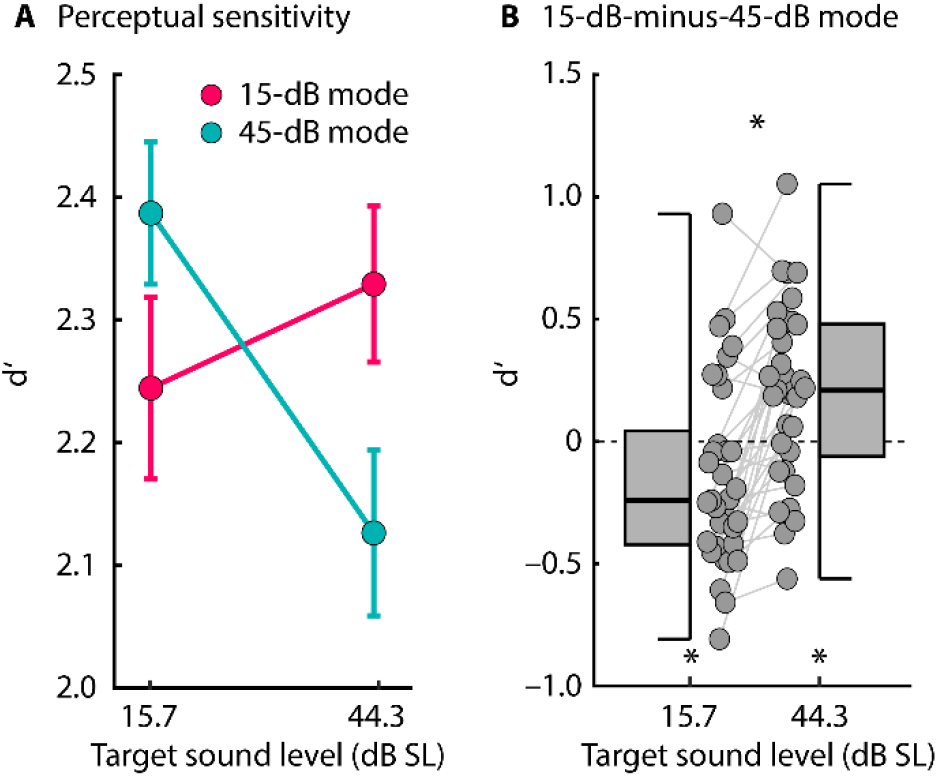
Perceptual sensitivity (d’) for detecting amplitude-modulated target noises in two sound-level contexts. **A:** d’ for detecting amplitude-modulated target bursts for different target-intensity categories (one centered on 15.7 dB SL and one on 44.3 dB SL) for the 15-dB SL and the 45-dB SL contexts. Error bars reflect the standard error of the mean (removal of between-subject variance^42^). The interaction is significant (p < 0.05). **B:** Difference in d’ between the 15-dB SL and 45-dB SL contexts, separately for low- and high-intensity targets. A positive value means a larger d’ for the 15-dB SL compared to the 45-dB SL context, and a negative value a larger d’ for the 45-dB SL compared to the 15-dB SL context. *p ≤ 0.05

Performance on the task was good, with all conditions exceeding a mean d’ value of 2. An rmANOVA indicated that the two factors Context and Target-Intensity Category exhibited a significant interaction (F_1,32_ = 34.610, p = 1.5×10^−6^, η^2^_p_ = 0.520; Figure 7A). Comparing d’ values between the two contexts, separately for each target-intensity category (lower-intensity targets centered on 15.7 dB SL; higher-intensity targets centered on 44.3 dB SL), revealed higher perceptual sensitivity to AM in the 15-dB context compared to the 45-dB context for high-intensity targets (t_32_ = 3.200, p = 0.003, r_e_ = 0. 492), but higher perceptual sensitivity to AM in the 45-dB context compared to the 15-dB context for low-intensity targets (t_32_ = −2.116, p = 0.042, r_e_ = 0.350; Figure 7B).

There was no effect of Context (F_1,32_ = 0. 0.266, p = 0.610, η^2^ = 0.008) nor an effect of Target-Intensity Category (F_1,32_ = 0. 587, p = 0. 449, η^2^_p_ = 0.018). The latter was expected because we normalized the AM depth for individual target-sound levels based on the Experiment II pre-test (Figure 6) to achieve equal task difficulty (i.e., AM-detection performance) for each target-sound level.

In sum, the results of Experiment II show that the detection of amplitude fluctuations (AM) in sounds is affected by sound-level statistical contexts. Consistent with a prediction-error account, sensitivity was enhanced for AM on higher-intensity targets compared to lower-intensity targets when ambient sound level was low, but that sensitivity was enhanced for AM on lower-intensity targets compared to higher-intensity targets when ambient sound level was high. The statistical results of Experiment II resemble the dependency of neural data on statistical context for the slow negative potential demonstrated in Experiment I (Figure 5), but does not mirror the context effects related to neural adaptation to sound-level statistics (Figure 4)^10,11,17,18,21^.

## Discussion

The current study investigated the effects of sound-level statistical context on neural activity and perceptual acuity for sound level. Sound-level statistics were manipulated by presenting white-noise bursts at levels drawn from one of two distributions with different modes (15 dB vs. 45 dB). Neural responses were affected by the statistical sound-level context in two different ways concurrently, providing evidence for both dynamic-range adaptation and prediction-error accounts (Experiment I). Perceptual sensitivity to amplitude modulation was affected by the statistical sound-level context in a way consistent with prediction-error processing, but not with neural adaptation (Experiment II).

In Experiment I, neural P1-N1 and P2-N1 amplitudes were sensitive to target intensity and to sound-level statistics. We observed a large effect of context (15-dB modal vs. 45-dB modal sound level) on higher-level target sounds, and much less of a context effect on lower-level sounds (Figure 4). P1-N1 amplitudes for lower-level target sounds showed a trend towards a reverse context effect (i.e., larger response in the 45-dB than 15-dB context), but this observation may need to be treated with caution due to the relatively small magnitude of the P1-N1 response (see Figure 4A). Our data, particularly P2-N1 amplitude effects, are consistent with the hypothesis that neurons in the auditory system adjust their sensitivity by shifting the dynamic response range depending on the modal sound level in an acoustic environment (schematically depicted in Figure 1B)^10,11,17,18,21^. These results are also consistent with observations from stimulus-specific adaptation, a related context-dependent form of adaptation^32,47–49^, showing enhanced sensitivity to sound level specifically for higher-level target sounds in lower-level compared to higher-level ambient contexts^32,50^. Here we used white-noise bursts, unlike in our previous work, in which we used pure tones^21^. White-noise bursts activate the full extent of the cochlear partition and associated auditory nerve fibers - this avoids the issue that pure tones with more intense levels recruit more off-frequency (i.e., non-best frequency) fibers, which, in turn, may contribute to effects of sound-level context using pure tones.

Neural adaptation to sound-level statistics has been observed at different stages along the auditory hierarchy, including the auditory nerve^18,19^, inferior colliculus^10,11,20,31^, and auditory cortex^51^. Until recently, investigations of adaptation to sound-level statistics were limited to animal models. The current data, together with recent magnetoencephalography data^21^, demonstrate response patterns that are consistent with adaptation to sound-level statistics in human auditory cortex. The mechanisms underlying statistical adaptation are not fully understood, but research in animals suggests that neural inhibition is likely an important contributor^4,17,52–55^.

Experiment I also revealed an increased slow negative potential that appeared to reflect prediction error. The negative potential was larger for higher-level target sounds in the 15-dB compared to the 45-dB context, and larger for lower-level target sounds in the 45-dB compared to the 15-dB context (shown as negative and positive deflections in the 15-dB context minus 45-dB context difference in Figure 5, respectively). This work is consistent with a large body of work on a deviant-related negative potential - the mismatch negativity^23–26,56^ - that has been linked to prediction-error processing (schematically depicted in Figure 1C)^27–30^: A sound whose features deviate from a previously established, predictable pattern elicit a negative potential originating in auditory cortices^57^. Experiment I shows that two processes - neural adaptation and prediction-error signals - can occur concurrently in auditory brain regions.

In Experiment II, we utilized the experimental paradigm (Figure 2) used to demonstrate that sound-level context influences neural responses to investigate effects on perception of amplitude modulation in sound. We observed that AM sensitivity for higher-intensity target sounds was higher in the 15-dB compared to the 45-dB context, but that AM sensitivity for lower-intensity target sounds was higher in the 45-dB compared to the 15-dB context (Figure 7). The behavioral effect of sound-level context resembled the effect of context on the EEG-recoded slow negative potential (Experiment I; Figure 5) and is consistent with prediction-error processing (Figure 1C). Our behavioral results are further in line with previous work investigating the effect of sound-level statistical context on sensitivity to sound level using pairs of noise bursts and longer intervals to allow for a behavioral response between noise pairs^33^. The current results and the results from this previous study suggest that prediction-error processes operate on shorter and longer time scales. Note that, in contrast to our neural data (Figure 4), there was no effect of target-intensity category on perceptual sensitivity (d’), because AM depth was normalized across target-sound levels to achieve equal AM-detection difficulty at different target levels. The effect of sound-level context on perceptual sensitivity is consistent with the hypothesis that prediction-error signaling optimizes perception in different statistical contexts^13,27–30,58^.

In the current study, noise bursts were presented every 0.5 seconds (with a 0.4-s silent interval), and targets were identical between contexts as were the noise bursts preceding the targets. Thus, 0.9 s would have elapsed between the end of the last filler (which could differ systematically between contexts) and the onset of the measured target. This is too long an interval for masking to be operating^59,60,61^, or for any other simple explanation of the effect. The differences in the effect of target intensity on AM-detection sensitivity must be due to the longer-term sound-level statistics, likely over multiple seconds^15^.

We investigated how sound-level statistics affect neural responses and perception by utilizing a paradigm in which white-noise bursts were presented in sound-level contexts with different modes (low and high). We show that neural-response sensitivity to sound level is affected by sound-level contexts in two different ways concurrently. First, the pattern of responses is consistent with the hypothesis that neural responsiveness changes with the modal ambient sound level, such that neurons are more sensitive and more responsive when ambient sound levels are low compared to when ambient sound levels are moderate to high (schematically depicted in Figure 1B). Second, the slow negative potential that occurs concurrently with these adaptation effects is consistent with the prediction-error hypothesis (schematically depicted in Figure 1C), such that neurons are most sensitive to sound levels that deviate from an established, predictable sound-level context. Perceptual sensitivity to sound-level modulation was also affected by sound-level statistical contexts in a way consistent with the prediction-error hypothesis: Sensitivity was highest for sounds whose levels deviated from the mode of the sound-level context. The data thus indicate that the auditory system uses statistical information about sound level to fine-tune perception dynamically by increasing sensitivity to low-probability events in a given statistical context. Adaptation to sound-level statistics was not apparently related to perceptual sensitivity.

## Methods and Materials

### Participants

Twenty-five adults participated in Experiment I (median age: 20 years; range: 18-31 years; 15 females), six adults participated in the Experiment II pre-test (median age: 23 years; range: 20-27 years; 4 females), and thirty-three adults participated in Experiment II (median age: 20 years; range: 17-31 years; 20 females). Two additional participants were recorded for Experiment I, but the data were excluded because no clear components in the event-related potential could be identified. One additional participant was excluded from Experiment II, because the person did not complete all blocks.

Participants reported no neurological disease or hearing impairment. They gave written informed consent prior to the experiment and were paid $5 CAD per half hour for their participation. The study was conducted in accordance with the Declaration of Helsinki, the Canadian Tri-Council Policy Statement on Ethical Conduct for Research Involving Humans (TCPS2-2014), and approved by the local Nonmedical Research Ethics Board of the University of Western Ontario (protocol ID: 106570).

### Acoustic stimulation

All experimental procedures were carried out in a single-walled sound-attenuating booth within a larger quiet testing room in the Western Interdisciplinary Research Building (Brain & Mind Institute, Western University). Sounds were presented via Sennheiser (HD 25-SP II) headphones and a Steinberg UR22 (Steinberg Media Technologies) external sound card. Stimulation was controlled by a PC or Laptop (Windows 7, 64 bit) running Psychtoolbox in MATLAB software (MathWorks, Inc.).

#### Estimation of sensation level

Before each experiment, the sensation level (SL) was determined for each participant for a white noise stimulus using a method-of-limits procedure that we have described in detail in previous work^62,63^. Sounds presented to participants during the experiments were presented relative to the estimated sensation level (SL).

#### Stimulus design

The experimental paradigm closely mirrored our previous work^21^. Noise bursts were presented in two different types of stimulation blocks (contexts) that differed with respect to the sound-level distributions from which a noise burst’s sound level was drawn: for one context the modal sound level was 15 dB SL, for the other it was 45 dB SL (Figure 2). In order to investigate the effects of sound-level statistics on neural responses and perception, we eliminated the confounding effects of different acoustics by ensuring that the bursts for which data were analyzed, as well as the immediately preceding burst were identical across contexts (i.e., 15 dB SL vs. 45 dB SL). Participants listened to six (Experiment I) or four (Experiment II) blocks, half of them with a 15-dB modal sound level, and the other half with a 45-dB modal sound level.

In each 700-second long block, 1400 white-noise bursts, each with a duration of 0.1 s, were presented. The onset-to-onset interval was kept constant at 0.5 s. In each block, eight sound levels ranging from 5 dB SL to 55 dB SL (step size: 7.143 dB SL) were each presented 30 times, yielding 240 target noises per block. These stimuli were identical across all blocks (black dots in Figure 2, middle/right). The 240 noise bursts immediately preceding each of these targets were also fixed, such that for each of the 30 presentations of one of the eight sound levels, the sound level of the preceding noise took on one of 30 sound levels (range: 5 dB SL to 55 dB SL; step size: 1.724 dB SL) without replacement. Thus, the same 240 experiment pairs were presented in each block. The sound levels for the remaining 920 filler-noise bursts were chosen randomly (range: 5 dB SL to 55 dB SL; step size: 0.1 dB SL) depending on the sound-level context (Figure 2). For half the blocks, sound levels were randomly chosen from a sound-level distribution with a 15-dB high-probability region (red dots in Figure 2, middle). For the other half of the blocks, sound levels were randomly chosen from a sound-level distribution with a 45-dB high-probability region (teal dots in Figure 2, right). High-probability regions had a width of 10 dB centered on 15 dB SL or 45 dB SL (Figure 2, left). The experimental pairs and the fillers were randomly intermixed in each block and presented such that at least one filler occurred before and after each experimental pair. Sound-level context blocks alternated, and the starting sound-level context was counter-balanced across participants.

Analysis of the influence of sound-level context on neural responses (Experiment I) and perceptual sensitivity (Experiment II) focused on target bursts (i.e., the second noise burst in each experimental pair): these were identical across the two contexts (black dots in Figure 2, middle/right) as was the preceding noise burst^21^.

### Experiment I: Effect of sound-level statistical context on neural responses

#### EEG recording and preprocessing

EEG signals were recorded at a 1024-Hz sampling rate from 16 electrodes (Ag/Ag-Cl-electrodes; 10-20 placement) and additionally from the left and right mastoids (BioSemi, Amsterdam, The Netherlands; 208 Hz low-pass filter). Electrodes were referenced to a monopolar reference feedback loop connecting a driven passive sensor and a common-mode active sensor, both located posterior on the scalp.

Offline data analysis was carried out using MATLAB software. Line noise (60 Hz) was suppressed in the raw data using an elliptic filter. Data were re-referenced to the average mastoids, high-pass filtered at a cutoff of 0.7 Hz (2449 points, Hann window), and low-pass filtered at a cutoff of 22 Hz (211 points, Kaiser window). Independent components analysis (runica method^64^; logistic infomax algorithm^65^; Fieldtrip implementation^66^) was used to identify and suppress activity related to blinks and horizontal eye movements. Data were divided into epochs ranging from −0.1 to 0.4 s (time-locked to burst onset). Epochs that exceeded a signal change of more than 180 μV for any electrode were excluded from analyses. All epochs were baseline-corrected by subtracting the mean signal in the 0.1 s prior to the noise-burst onset from each time point of the epoch (separately for each channel).

#### Data analysis

##### P1-N1 and P2-N1 peak-to-peak amplitude

Data analysis focused on the P1-N1 and P2-N1 peak-to-peak amplitudes, as in previous work^34–38^. Estimation of P1, N1, and P2 peak amplitudes for each participant can be challenging when responses are strongly adapted as in the current experiment. In order to overcome this challenge and estimate individual responses amplitudes of the P1, N1, and P2 components of the event-related potentials, we utilized a simplified jackknifing approach that retrieves amplitude estimates for each participant^67^. For a specific condition, data from one participant were left out, while the response time courses (averaged across a fronto-central electrode cluster) of the remaining participants were averaged. P1, N1, and P2 amplitudes were extracted from this leave-one-out average as the mean signal within a 10-ms time window centered on the P1, N1, and P2 peaks, respectively. This procedure was repeated such that data from each participant were left out once. In order to obtain the participant-specific estimate using the simplified jackknifing approach, we applied the following equation^67^:

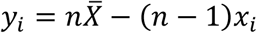

where *y*_*i*_ refers to a participant-specific amplitude estimate of one component of the event-related potential; *x*_*i*_ to the sub-average amplitude for which the participant data were left out, *n* to the number of participants, and 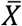 to the mean across sub-averages in *x*. Critically, although jackknifing makes uses of sub-averages across participants to calculate stable estimates of individual data, the approach effectively transfers the subaverage values back to the individual participant so that they can be interpreted similarly to those derived from original single-participant data^67^.

##### Response sensitivity to sound level

In order to investigate how sound level affects response amplitude (across statistical contexts), trials from all context blocks were sorted into one of five categories according to their sound level, ranging from 5 to 55 dB (width 20 dB). Single-trial response time courses were averaged separately for each sound-level category. For each participant, the P1-N1 and the P2-N1 amplitude difference was extracted from the averaged time courses, separately for each sound-level category. In order to investigate whether responses increased with increasing sound level, a linear function was fit to the amplitude as a function of sound level, separately for P1-N1 and P2-N1 amplitudes. The slope of the linear function indicates the degree of response sensitivity to sound level. Each participant’s data yielded one slope value for the P1-N1 and the P2-N1: we tested whether the slopes differed from zero using a one-sample t-test.

##### Context-dependent response sensitivity

For this analysis, only responses to target bursts with the 8 different critical sound levels were analyzed (black dots in Figure 2). The number of trials at each of the 8 sound levels in the two contexts was relatively low (90 trials) and the response magnitudes were relatively small due to strong neural adaptation. In order to increase the number of trials in the response average, thereby increasing the signal-to-noise ratio, we binned the target responses into two categories based on target sound levels (separately for the two contexts). After baseline correction, single-trial time courses for noise bursts with a level below 30 dB SL (i.e., four levels between 5 and 26.4 dB with a mean of 15.7 dB SL) were averaged as were the time courses for bursts with a level above 30 dB SL (i.e., four levels between 33.6 and 55 dB with a mean of 44.3 dB SL). P1-N1 and P2-N1 response magnitudes were analyzed using a repeated-measures ANOVA with the factors Context (15 dB SL, 45 dB SL) and Target-Intensity Category (2 levels, centered on 15.7 and 44.3 dB SL).

### Experiment II pre-test: Determination of AM-detection thresholds at different sound levels

A psychophysical pretest was conducted using the method of constant stimuli, in order to estimate the AM depth corresponding to the detection threshold for a 50-Hz AM in a 0.1-s duration white-noise burst, for different sound levels. The slope of the function relating intensity to AM-detection sensitivity was then used in Experiment II (Figure 6).

White-noise bursts (0.1 s duration) were presented every 0.5 s. Targets bursts were amplitude modulated at a rate of 50 Hz; fillers were unmodulated. Participants listened to three blocks of noise bursts, each containing 160 targets and 906 fillers. Participants were instructed to press a button on a keyboard as soon as they detected a target. Each target was presented at one of four sound levels (i.e., 15, 25, 35, or 45 dB SL) and one of eight AM depths (from 2% to 100%, in steps of 14%). In each block, five targets were presented at each sound-level and AM-depth combination. The sound levels for fillers were randomly drawn from a uniform distribution ranging from 5 dB SL to 55 dB SL. Between three and eight fillers were interposed between successive targets. Participants performed one to three brief training blocks to familiarize them with the stimulation and task.

A target was considered detected if a key press was made between 0.1-1.1 s after the target onset. The proportion of detected targets was calculated for each sound level and AM depth, pooled across all six participants. A logistic function was fit to the proportion of detected targets as a function AM depth, separately for the four different sound levels. For each sound level, the AM depth that yielded 50% detected targets was estimated from the fitted logistic function. In order to estimate the AM depth for any of the eight target-sound levels used in Experiment II (Figure 2), we fitted a linear function to the 50% thresholds as a function of the four pre-test target levels. The slope from the linear fit was used in Experiment II to relate AM depth to target-sound level.

### Experiment II: Effect of sound-level statistical context on AM detection

The stimuli of Experiment II were identical to those in Experiment I, except that target bursts (see Figure 2; black dots) were amplitude modulated at 50 Hz while fillers remained unmodulated (Figure 6). Participants performed the AM-detection task, as in the Experiment II pre-test, for four stimulation blocks. Sound levels in the blocks were drawn from one of the two sound-level distributions (with a mode of 15 dB SL or 45 dB SL) as in Experiment I. Blocks alternated between the 15 dB SL context and the 45 dB SL context (counterbalanced across participants). As in Experiment I, the sound levels of targets, and the level of preceding bursts, were identical across contexts (Figure 2, black dots). Hence, any difference in AM-detection performance between contexts can be attributed to the sound-level statistics.

For Experiment II, we wished to equalize the AM-detection rate across target-sound levels in order to avoid differences in task difficulty for different target intensities. This involved estimating the AM depth that would yield 75% detected targets at one sound level (40 dB SL) in a pre-experiment block and calculating the AM depth for all target intensities relative to this 75%-AM depth using the estimated slope from the Experiment II pre-test that relates AM-detection thresholds to sound levels. In the pre-experiment block, ten 40-dB SL targets at each of eight modulation depths (2% to 100%, in steps of 14%) were presented among 450 fillers drawn from a uniform level distribution that ranged from 5 to 55 dB SL. The proportion of detected targets was calculated for each AM depth and a logistic function was fit to the data to obtain the AM depth that yielded a 75% detection rate at 40 dB SL. AM depths for all target intensities were then estimated from the slope of the Experiment II pre-test relative to the AM depth yielding the 75% detection rate at 40 dB SL. Note that the identical AM depths were used for targets in both statistical contexts during the main part of Experiment II, and any difference in AM-detection performance between contexts can thus be attributed to the sound-level statistics.

For the analysis of Experiment II data, a target was considered detected if a key press was made within 0.1-1.1 s after the target onset. As in Experiment I, we binned targets into two categories based on burst intensity in order to increase the number of trials (and thus signal-to-noise ratio). Targets with a sound level below 30 dB SL (mean of 15.7 dB SL) were analyzed separately from those with a sound level above 30 dB SL (mean of 44.3 dB SL). For each participant, the proportion of detected targets was calculated separately for each sound-level context (15 dB SL, 45 dB SL), and each target-intensity category (15.7 dB, 44.3 dB). In order to calculate perceptual sensitivity (d’), the false alarm rate was calculated using a method applicable to paradigms with high event rates such as the current one^68^. Perceptual sensitivity (d’) values were subjected to a repeated-measures ANOVA with the factors Context (15 dB SL, 45 dB SL) and Target-Intensity Category (15.7, 44.3 dB SL).

### Effect sizes

Throughout the manuscript, effect sizes are provided as partial eta squared (η_p_^2^) for repeated-measures ANOVAs and as r_e_ (r_equivalent_) for t-tests^69^.

## Acknowledgements

Research was supported by the Canadian Institutes of Health Research (MOP133450 to I.S. Johnsrude). BH was supported by a BrainsCAN Tier I postdoctoral fellowship (Canada First Research Excellence Fund; CFREF) and by the Canada Research Chair program. We thank Kurdo Araz for his help during data collection.

## Author contributions

The experiments were performed in the Psychology Department and Brain and Mind Institute at the University of Western Ontario, London, Canada. BH, TA, ISJ designed the study. BH and TA programmed the experiments, collected the data, and analyzed the data. BH and ISJ wrote the manuscript. All authors reviewed and revised the manuscript.

## Data availability

The datasets generated during the current study are not publicly available because participant consent for public-data sharing was not obtained. Data are available from the corresponding author on reasonable request.

